# Genomic signatures of the unjamming transition in compressed human bronchial epithelial cells

**DOI:** 10.1101/2020.09.01.277962

**Authors:** Margherita De Marzio, Ayşe Kılıç, Enrico Maiorino, Jennifer Mitchel, Maureen McGill, Robert Chase, Jeffrey J. Fredberg, Jin-Ah Park, Kimberly Glass, Scott T. Weiss

## Abstract

Epithelial tissue has the capacity to switch from a collective phase that is quiescent, solidlike and non-migratory to one that is dynamic, fluid-like and migratory. In certain physiological and pathophysiological contexts this phenotypic switch has been attributed not to the well-known epithelial-to-mesenchymal transition, EMT, but rather to the recently discovered unjamming transition, UJT. UJT has been characterized thus far mainly at functional and morphological levels whereas underlying genome-wide molecular events remain largely unexplored. Using primary human bronchial epithelial cells and one well-defined trigger of UJT –mechanical compression– here we combine temporal RNA-Seq data and Protein-Protein Interaction networks to provide the first genome-wide analysis of UJT. Our results show that compression induces a multiphasic transcriptional response characterized by an early activation of genes regulating the membrane and actomyosin structure, and a delayed activation of genes regulating the extracellular matrix and cellmatrix interactions. This biphasic response is mediated by a cascade of signaling processes that promotes actin polymerization through the recruitment of integrin-ECM adhesive complexes and promotes increased cellular motility through activation of AP-1 transcription factors via ERK and JNK pathways. These findings, taken together, show that the UJT program is not the result of any single signaling pathway but rather comprises a coordinated interplay of downstream pathways including development, fate selection, energy metabolism, cytoskeletal reorganization, and adhesive interaction with extracellular matrix.

## Introduction

While performing its routine barrier and immune functions, the cellular collective that defines a confluent epithelial tissue is typically quiescent, solid-like, and non-migratory. In a variety of circumstances, however, the epithelial collective undergoes an unjamming transition (UJT) to become dynamic, fluid-like, and migratory.^1-14^ For example, experimental data from both *in vitro*^1-3,5,7,11,15^ and *in vivo*^2,6^ studies have shown that UJT occurs spontaneously during physiological events such as embryonic development^2,6,9^ as well as during pathophysiological events including wound repair^1,16^ and cancer metastasis^4,8,9,17,18^. In addition, stimuli such as radiation^19^, stretch^2^, and mechanical compression^2,11,15^, all of which are associated with lung diseases, also have the capacity to trigger UJT. In the jammed phase each cell becomes virtually frozen in place –trapped by its immediate neighbors– in a collective phase where intercellular rearrangements are rare. Conversely, in the unjammed phase intercellular rearrangements are frequent and the confluent cellular collective moves cooperatively, collectively and vigorously in packs and swirls reminiscent of fluid flow^1,20^.

Experimental studies to date have emphasized functional, morphological and, to only a limited extent, molecular features that define the UJT^2,9,11,17^ and distinguish it from the canonical epithelial-to-mesenchymal transition (EMT)^15^. After EMT, for example, cell-cell junctions become degraded, barrier function becomes compromised, apico-basal polarity is lost, and mesenchymal markers appear. After UJT, by contrast, none of these events pertain^15^. Rather, cell shapes become more and more variable, and these morphological changes coincide with increased motility and cellular cooperativity^11,15^. As such, during UJT the cell layer becomes migratory while retaining its full epithelial character.

Recent insights have begun to elucidate underlying molecular events. In a breast cancer model system, for example, *RAB5A* triggers unjamming by promoting internalization of *EGFR* leading to hyperactivation of the kinase ERK1/2 and phosphorylation of the actin nucleator *WAVE2*^17^. In well-differentiated primary human bronchial epithelial cells (HBECs), mechanical compression unjams the epithelial layer^11,15^ and activates signaling pathways including EGFR^21-23^, TGF-β receptor^24-27^ and ERK^15,23^. In the same model system, it also induces procoagulant factors^28,29^ and induces an asthma-like transcriptional pattern that involves inflammatory, fibrotic and remodeling processes^30^.

These results establish UJT as a complex multifactorial program. In doing so they also emphasize the necessity of obtaining a more comprehensive molecular assessment, but such an assessment is difficult to achieve with single-gene targeted approaches. The development of RNA-Sequencing (RNA-Seq) technologies^31^ now offers an opportunity to profile the genome-wide transcriptomic signature of UJT. RNA-Seq measures the total cellular content of RNA at the transcript-level, providing information on the key genes activated in a specific condition. This information can be further expanded by integrating molecular interaction data^32-34^, which can be used to infer the functional and causal relationships behind a genomic pattern^35^.

Here we report the first comprehensive genomic profiling of epithelial unjamming. We do so by performing RNA-Seq of well-differentiated HBECs after compression, which is a known trigger of the UJT. We analyze the transcriptional patterns of compressed HBECs over three time points (baseline, 3 hours post-compression and 24 hours post-compression) and integrate the RNA-Seq data with protein-protein interaction (PPI) networks to identify the molecular pathways inducing these patterns. Our analysis shows that compression-induced UJT is not the result of a single biological process but rather the result of the coordinated interplay of multiple downstream pathways involving actin repolymerization, ERK and JNK signaling, and ECM reorganization.

## Results

### Compression-induced UJT activates a bimodal transcriptional response

To investigate the biological mechanisms driving the UJT, we harvested HBECs, grew them to confluence under Air Liquid Interface (ALI), and then exposed them to an apico-to-basal pressure difference of 30 cm H_2_O^15,26,29,30,36-38^ (see Materials and Methods). Following the same experimental protocol described in Kılıç et al.^30^, this pressure was maintained for 3 hours and then released (Fig. 1a). We assessed the gene expression profile of compressed HBECs using bulk RNA-Seq at two time points: immediately after pressure release (3-hour time point) and when cells had rested for additional 21 hours after pressure removal (24-hour time point). For both time points, RNA-Seq was also performed on non-compressed control HBECs.

**Fig. 1.**
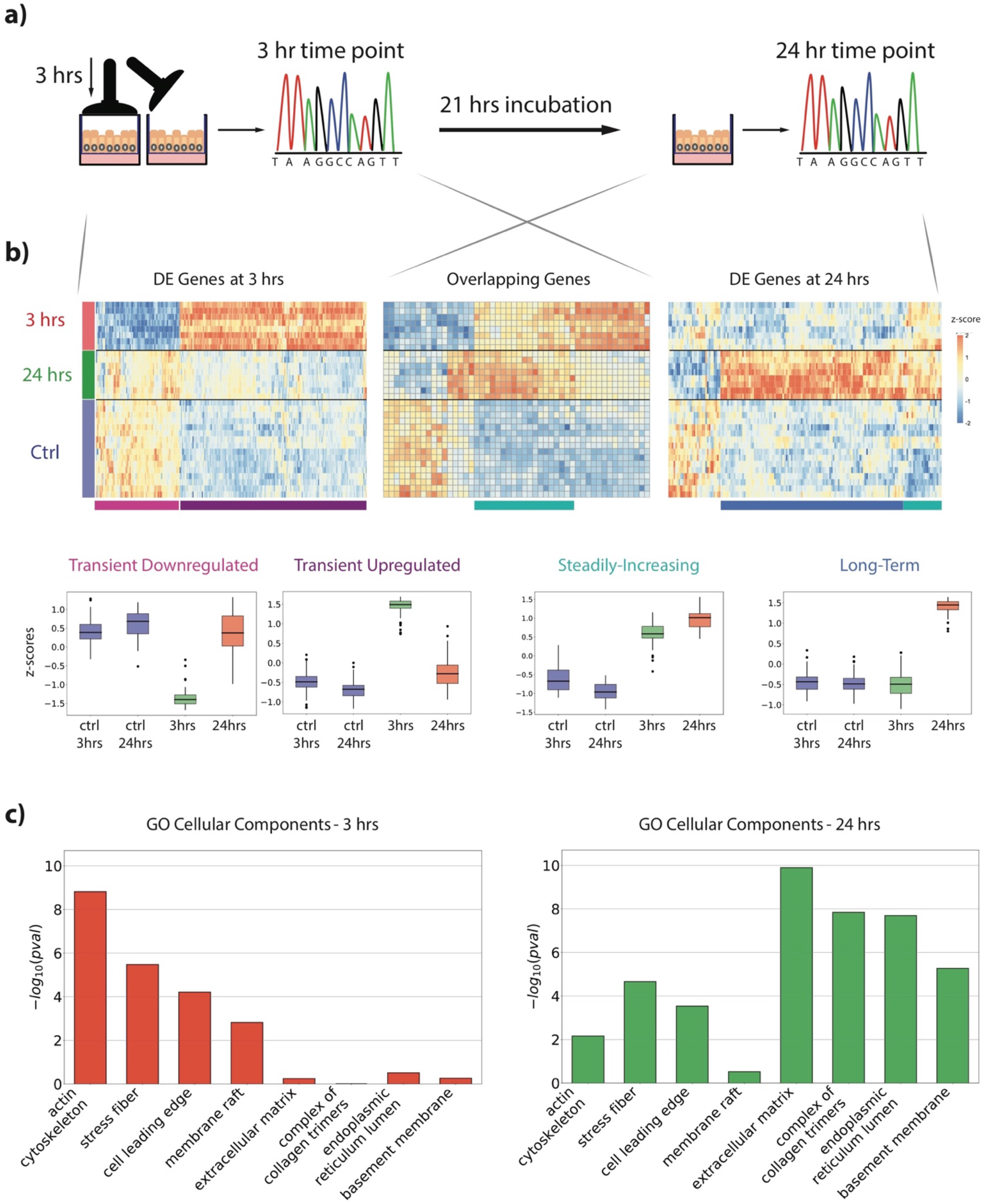
Overall scheme of the transcriptional patterns activated by compression. **a** The experimental setup: mechanical compression was applied to well-differentiated HBECs for three hours and subsequently released. RNA-Seq was extracted from post-compression HBECs immediately after pressure release (3-hour time point) as well as after an additional 21 hours (24-hour time point). **b** (Top) Heatmaps of the normalized expression values at each time point (control, 3 hours, 24 hours) for genes DE at 3 hours (left), genes DE at both 3 hours and 24 hours (center), and genes DE at 24 hours (right). Rows are z-score normalized for visualization purposes. **b** (Bottom) Boxplots of the genes’ z-scores in the three transcriptional regimes identified by clustering the data shown on top: transient downregulated, transient upregulated, steadily-increasing, and long-term regimes. Each transcriptional cluster is pointed out at the bottom of the heatmap through a specific color. **c** Log(p-value) of a merged subset of GO Cellular Components enriched at 3 hours and 24 hours. At 3 hours (left), terms associated with the cytoskeleton and cellular membrane are highly enriched compared to the terms associated with the extracellular space. At 24 hours (right), this trend is inverted, and GO terms related to collagen complexes and extracellular matrix become highly overrepresented.

Recent studies^11,15^ investigated the morphological features of post-compression HBECs under the experimental setup described above, and showed that compressed HBECs exhibit all the cardinal features of the UJT^39,40^. Beginning at 8 hours post-compression, cells become more elongated and variable in shape^11,15^ and the epithelial layer becomes more fluid-like, displaying swirling patterns of migratory behavior^11,15^. At 24 hours post-compression, HBECs show structural and dynamic signatures of an unjammed epithelium^11,15^ and show progressively increasing migration and elongation up to 72 hours post-compression^15^.

In parallel to these morphological changes, our RNA-Seq data showed that compression activated a multiphasic transcriptional response. We assessed this response by performing differential expression analysis of post-compression HBECs at 3 hours and 24 hours with respect to control samples at the same time points.

At the 3-hour time point, differential expression analysis identified 283 differentially expressed (DE) genes (fold-change (FC)>1.5 and False Discovery Rate (FDR)-adjusted p-value<0.05); this included 89 downregulated and 194 upregulated genes (Supplementary Table 1). At the 24-hour time point, we observed an additional wave in the transcriptional activity of cells with 42 genes under-expressed and 171 genes overexpressed (Supplementary Table 2).

DE genes at 3 hours versus 24 hours revealed striking differences in the post-compression transcriptional responses. To characterize these differences, we computed the z-scores of the normalized expression values and performed hierarchical clustering on the DE genes identified at each time point (see Materials and Methods). Based on the clustering results, we considered the three main classes of genes that exhibited characteristic transcriptional responses. A significant portion of the DE genes at 3 hours (233 out of 283, 82%) returned their expression to baseline level at 24 hours (Fig. 1b, Supplementary Table 3a, b). As a quantitative confirmation of this observation, the Pearson correlation coefficient of the FCs between 3 hours versus control and 24 hours versus 3 hours was *ρ*= −0.84. These transcriptional changes suggest the activation of a *transient* regime which reflects the immediate, but temporary, response of cells to the mechanical stimulus.

For a subset of 109 DE genes at 24 hours (Supplementary Table 4), transcriptional activity was not affected at 3 hours but increased only at 24 hours (Fig. 1b). We retrieved this behavior independently on the control time point used for comparison, excluding possible biases due to changes over time in the expression profile of control samples. This result indicates that the mechanical stimulus caused a delayed effect on HBEC gene expression and activated a specific, *long-term* transcriptional program. This transcriptionally active program at 24 hours corresponds to the beginning of a large and sustained increase in cellular unjamming which continues to at least 72 hours^15^.

Finally, 42 genes (Supplementary Table 5), that included both overlapping genes and DE genes at 24 hours, exhibited a stable increase in their expression levels (Fig. 1b). This reflects activation of a *steadily-increasing* transcriptional regime where the alterations generated by compression are positively reinforced with time.

Overall, compression triggered a dynamic response where transient effects at 3 hours were accompanied by both steadily increasing and delayed effects unfolding over time. Since HBECs begin to unjam several hours after pressure release^15^, we hypothesized that the first transient perturbation triggered a cascade of signaling events that induced a secondary transcriptional response detected at 24 hours. The outcomes of the associated downstream processes were the key molecular components driving and sustaining cell unjamming.

In order to test this hypothesis, we performed a Pathway Overrepresentation test on cellular component (CC) and biological process (BP) Gene Ontology (GO) annotations^41,42^ using the R package Cluster Profiler^43^; results were selected based on an FDR-corrected p-value threshold of 0.05 (see Materials and Methods).

Enrichment analysis using CC annotations revealed that at 3 hours DE gene products were localized mainly in the cell membrane and cytoplasm (Fig. 1c), with actin cytoskeleton (p-adj. = 1.6e-09) and cell leading edge (p-adj. = 6.3e-05) among the most enriched components. At 24 hours, the effects of the mechanical stress propagated from the cellular surface towards the endoplasmic reticulum and the extracellular matrix space, exhibiting high enrichment in the endoplasmic reticulum lumen (p-adj. = 2.1e-08), complex of collagen trimers (p-adj. = 1.4e-08), and basement membrane (p-adj. = 5.5e-06). Additionally, the transient, steadily-increasing, and long-term regimes segregated into different GO biological processes, highlighting distinct functions for each gene subset.

We identified the major categories enriched in each response by visualizing enriched pathways through a network representation (Fig. 2). Transiently downregulated genes at 3 hours were exclusively annotated to developmental and differentiation pathways (Fig 2a and Supplementary Table 6a). Temporary suppression of cell fate decision and cell cycle functions in response to stress has been observed in several studies^44,45^ and this behavior is directly related to the capacity of cells to adapt to environmental conditions.

**Fig. 2.**
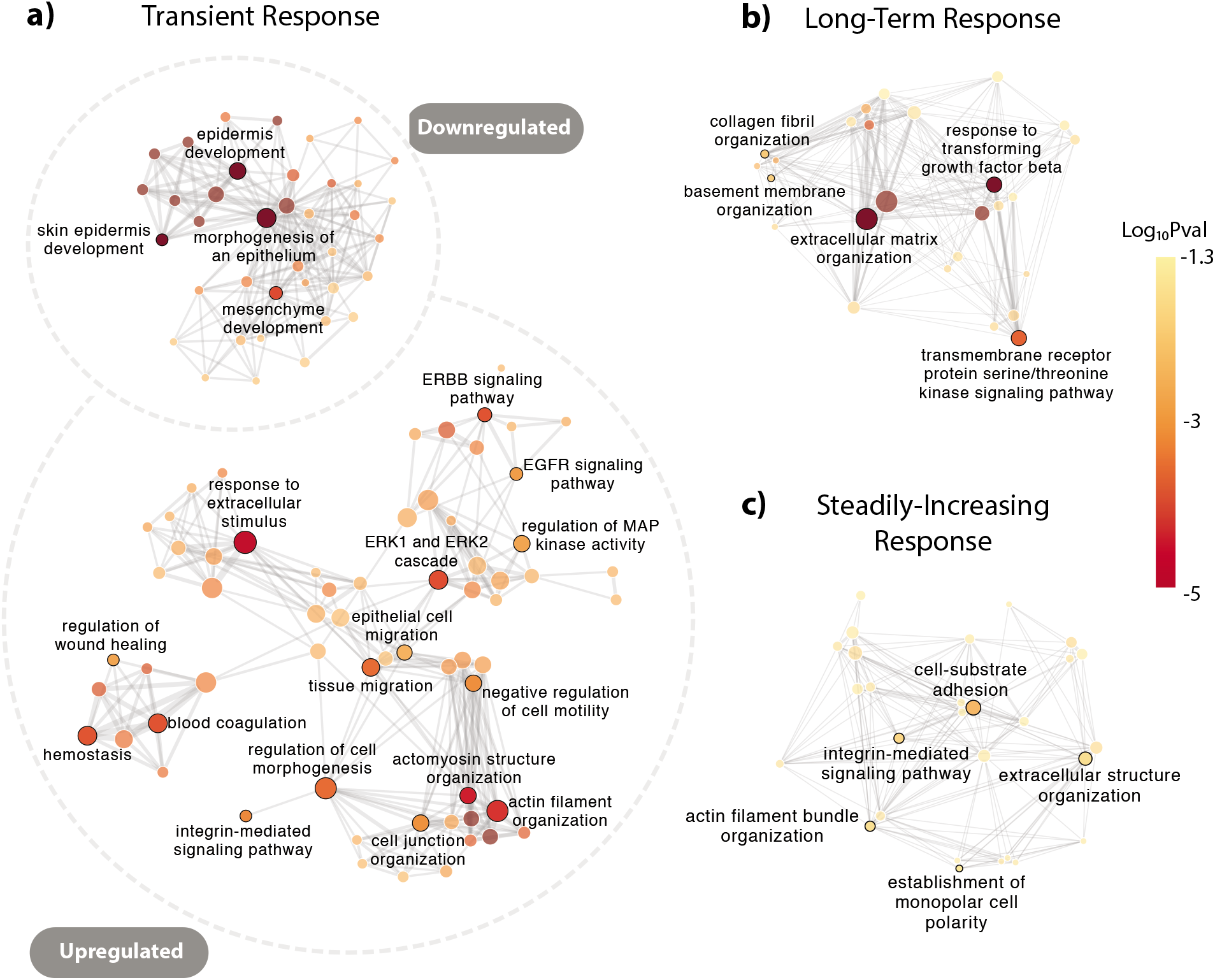
Pathways enriched in the three transcriptional regimes triggered by compression. For each transcriptional response, we visualized enriched GO Biological Processes (BP) using a network representation where each node represents a pathway and edges between pairs of nodes represent common DE genes annotated to both pathways. For visualization purposes we showed only edges with more than 3 common DE genes. Node size and color are based on the number of DE genes annotated to the pathway and its adjusted p-value, respectively. **a** GO BP enriched in the transient response, **b** GO BP enriched in the long-term response, **c** GO BP enriched in the steadily-increasing response.

Conversely, transiently overexpressed genes at 3 hours mapped to five main biological processes (Fig. 2a and Supplementary Table 6b): actomyosin structure organization (p-adj. = 1.7e-04), tissue migration (p-adj. = 8.1e-04), ERK1/2 cascade (p-adj. = 4.4e-04), regulation of MAP kinase activity (p-adj.=3.7e-03), and wound healing (p-adj. = 3.7e-03). Notably, several genes annotated to these pathways are known to be involved in the early activation and regulation of cell shape and movement.

DE genes annotated to cytoskeleton reorganization comprised the actin genes *ACTA1* and *ACTG1*, and actin-binding and regulating genes such as *CDC42EP2, RHOB, FHDC1, LCP1, PLS3*, and *RASSF1*. Additionally, genes annotated to cell migration included the soluble factors *HBEGF, F3*, and *SRF*, as well as genes involved in actin and myosin regulation such as *HMOX1, EGR3, MYH9*, and *MYADM*. These proteins are known to induce collective movement by promoting actin repolymerization and formation of focal adhesion complexes^46,47^. The enrichment analysis also highlighted multiple genes involved in the ERK cascade signaling, such as sprouty proteins *SPRED1, SPRED2*, and *SPRY2* and the FAK subfamily gene *PTK2B*.

Pathways associated with wound healing included angiogenesis, blood coagulation and platelet aggregation. These pathways are active during development and repair processes, and require reorganization of the cytoskeleton along the leading edge to either facilitate tissue growth or wound closure^48^.

While DE genes at 3 hours had a role in the earliest molecular events of cellular unjamming, genes involved in the long-term transcriptional response were mainly annotated to TGF-β receptor and extracellular matrix (ECM) signaling processes (Fig 2b and Supplementary Table 6c). Extracellular structure organization and collagen fibril organization were enriched in matrix metalloproteinases such as *MMP2* and *MMP10*, and in multiple collagen alpha chain genes, including collagens type-I, III, IV, and V. Overexpression of these genes points to an active reorganization of the ECM at 24 hours, a key process to maintain cell movement by allowing flexible attachment of cells to the substratum^48-50^. GO terms associated with transforming growth-factor receptor were annotated to target genes of TGF-β1-signaling such as *NOX4, THBS1* and *PMEPA1*.

Finally, enrichment analysis of the steadily-increasing genes revealed annotations to molecular processes involving cell-substrate adhesion, actin filament bundle organization, and extracellular structure organization (Fig. 2c and Supplementary Table 6d). Among these genes, TGF-β1-induced transcript 1, *TGFB1I1*, as well as actin-binding genes *FLNA, CNN1*, and *MYL9* showed fold-increases larger than 2 at both 3-hour and 24-hour time points. Increased co-expression of these proteins has been recently connected to the formation of ROCK-dependent actin fibers downstream of *TGFB1I1* induction^51^ suggesting a possible role of this gene in regulating cell migration^52,53^.

Our analysis highlights a transcriptional reprogramming of the active regulators of cell shape and motility at the 3-hour time point and of the active regulators of cell-substratum interactions at the 24-hour time point. In a confluent monolayer of MDCKII cells, it has been recently shown that the structural and dynamical changes of cells undergoing UJT are accompanied by increased glycolytic activity and larger mitochondrial membrane potential^54^. This work led us to investigate the potential role of metabolic processes in our model system. We found that GO terms enriched at 3 hours included regulation of lipid metabolic process (p-adj.=7e-03), regulation of small molecule metabolic process (p-adj.=8.5e-03), and regulation of cellular carbohydrate metabolic process (p-adj.=2.8e-02). These processes involve oxidization of fatty acids that are subsequently metabolized in mitochondria to produce ATP^55-58^. Genes annotated to the enriched pathways included known gluco-regulators and transporters^59-61^ such as *SNCA, SESN2*, and *SLC7A11* (Supplementary Table 7). The mitochondrial carrier protein *SLC25A25* was also highly upregulated at 3 hours (FC=4.9), suggesting increased mitochondrial activity in support of ATP production^62,63^. Notably, we didn’t observe any enrichment in metabolic processes at 24 hours. Our results hint at a potential correlation between cell energy metabolism and the UJT and emphasize the necessity of further experiments to elucidate the molecular mechanisms behind it.

### Integrins, ERK and JNK signaling pathways drive the epithelial cell response to compression

The dynamic evolution of the post-compression gene expression profile suggests activation of an underlying cascade of signaling events in the 21 hours following pressure release. To identify these processes, we first determined which pathways of the Kyoto Encyclopedia of Genes and Genome (KEGG)^64-66^ database were enriched in a merged set of DE genes at 3 hours and 24 hours (see Materials and Methods). We set the FC threshold of the differential expression analysis to 1.2 and the FDR-adjusted p-value threshold to 0.05, which resulted in a total of 1512 DE genes across the two time points.

Among the 29 statistically significant KEGG pathways (Supplementary Fig. 1), focal adhesion and MAPK signaling were the most enriched (FDR-adjusted p-values=4.74e-05 and 9.95e-05 respectively). Similar pathways emerged also using the Reactome database^67^ (Supplementary Table 8). Given the predominant role of focal adhesions in cytoskeleton reorganization and cell migration, we analyzed the DE genes associated with this pathway at each time point in greater detail. At the membrane localization, compression altered the expression of several ITGA and ITGB integrins (Fig. 3a). While *ITGB3* and *ITGB6* were overexpressed at both time points, other integrin chains were selectively induced at either 3 hours (*ITGA2* and *ITGA5*) or at 24 hours (*ITGA4, ITGB5*, and *ITGAV*), possibly adjusting to the changing ECM.

**Fig. 3.**
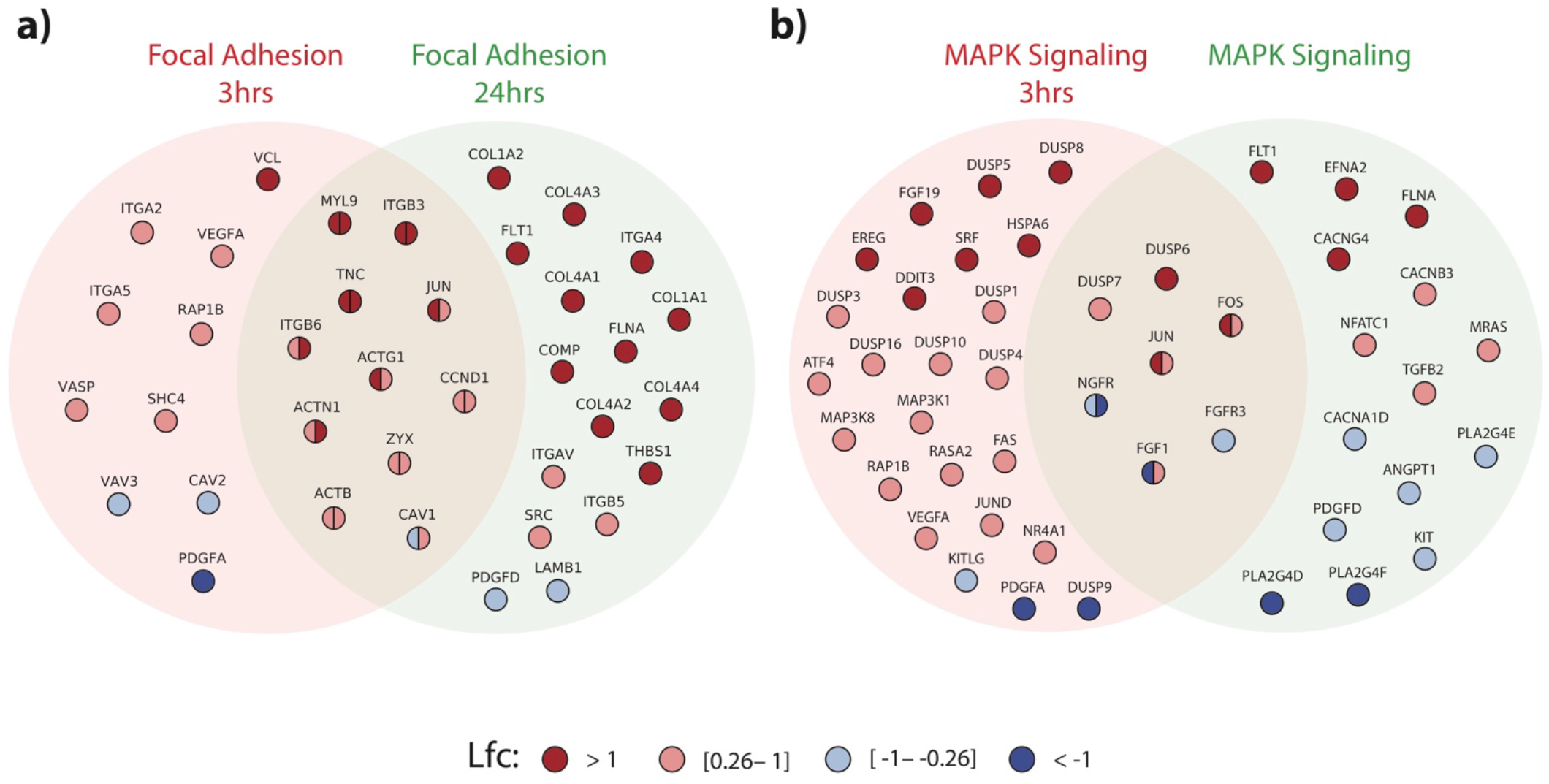
DE genes annotated to the KEGG focal adhesion and MAPK signaling pathways. Overview of DE genes at 3 hours and 24 hours, color-coded based on their log-fold-change, that are also annotated to the **a** focal adhesion, or **b** MAPK signaling pathways.

In addition to these transmembrane receptors, we found upregulation of the small GTPase protein encoded by *RAP1B* (3 hours) and of multiple actin-binding adaptor genes such as *VCL* (3 hours), *ZYX* (3 hours and 24 hours), *SRC* (24 hours), and *FLNA* (24 hours); these genes are known to transmit mechanical forces between the ECM and the actin cytoskeleton^68-71^. ECM proteins, including collagen chains and the tenascin *TNC*, were overexpressed only at 24 hours. These findings highlight the recruitment of several integrin-ECM adhesive complexes and suggest a reorganization of the actomyosin network downstream to the integrin signaling^68-71^.

The MAPK signaling pathway was also highly enriched. Multiple components involved in the ERK1/2 pathways were strongly overexpressed (Fig. 3b), including genes encoding growth factors, Ras-family proteins *RAP1B* (3 hours), *RASA2* (3 hours) and *mRAS* (24 hours), and the transcription factors *FOS* (3 and 24 hours), *ATF4* (3 hours), and *SRF* (3 hours). Compression also activated several molecules related to the JNK signaling pathways, such as *MAP3K1* (3 hours) and *MAP3K8* (3 hours), and the proto-oncogenes *JUND* (3 hours) and *JUN* (3 and 24 hours). These results, combined with the overexpression of genes in the DUSP family at 3 hours, which are known counterregulators of MAPK^72^, hint at a differential fine-tuning of MAPK signaling.

In contrast with the consistent overexpression of integrins, other growth factors and receptors, such as *FGFR3* (3 and 24 hours), *ANGPT1* (24 hours), and *PDGFA/D* (3 and 24 hours) were downregulated. Similarly, the expression of the cytosolic phospholipases *PLA2G4D, PLA2G4E* and *PLA2G4F*, were reduced at 24 hours.

To extract additional information on the active signaling cascade involving these focal adhesion and MAPK components, we combined our RNA-Seq data with known protein interactions contained in Protein-Protein Interaction (PPI) databases. PPI data can be represented as networks whose nodes are proteins that are linked to each other by physical binding interactions^73^. The PPI network represents a powerful tool to infer information on the cell-signaling pathways occurring in a specific process. In particular, analysis of network paths, defined by the sequences of edges that connect two nodes, can be used to identify the molecular processes that transmit signal between two proteins.

Following a procedure similar to the one proposed in ^74^, we modeled a signal transduction process as a sequential path on the network that starts from a receptor, connects the receptor to multiple kinases, and leads to a transcription factor (Fig. 4a). This Receptor-Kinase-Transcription Factor (RKT) path representation mimics the typical pattern of signaling cascades, where the signaling transmission is triggered by membrane-bound receptors, propagates via intracellular kinases, and ultimately regulates the activity of transcription factors^75^. We used the databases developed in ^76^ and ^77^ for the selection of the receptors and kinases respectively. To focus our analysis on the RKT paths that have a major impact on the long-term transcriptional response, we selected the transcription factors that target specifically DE genes at 24 hours (see Materials and Methods).

**Fig. 4.**
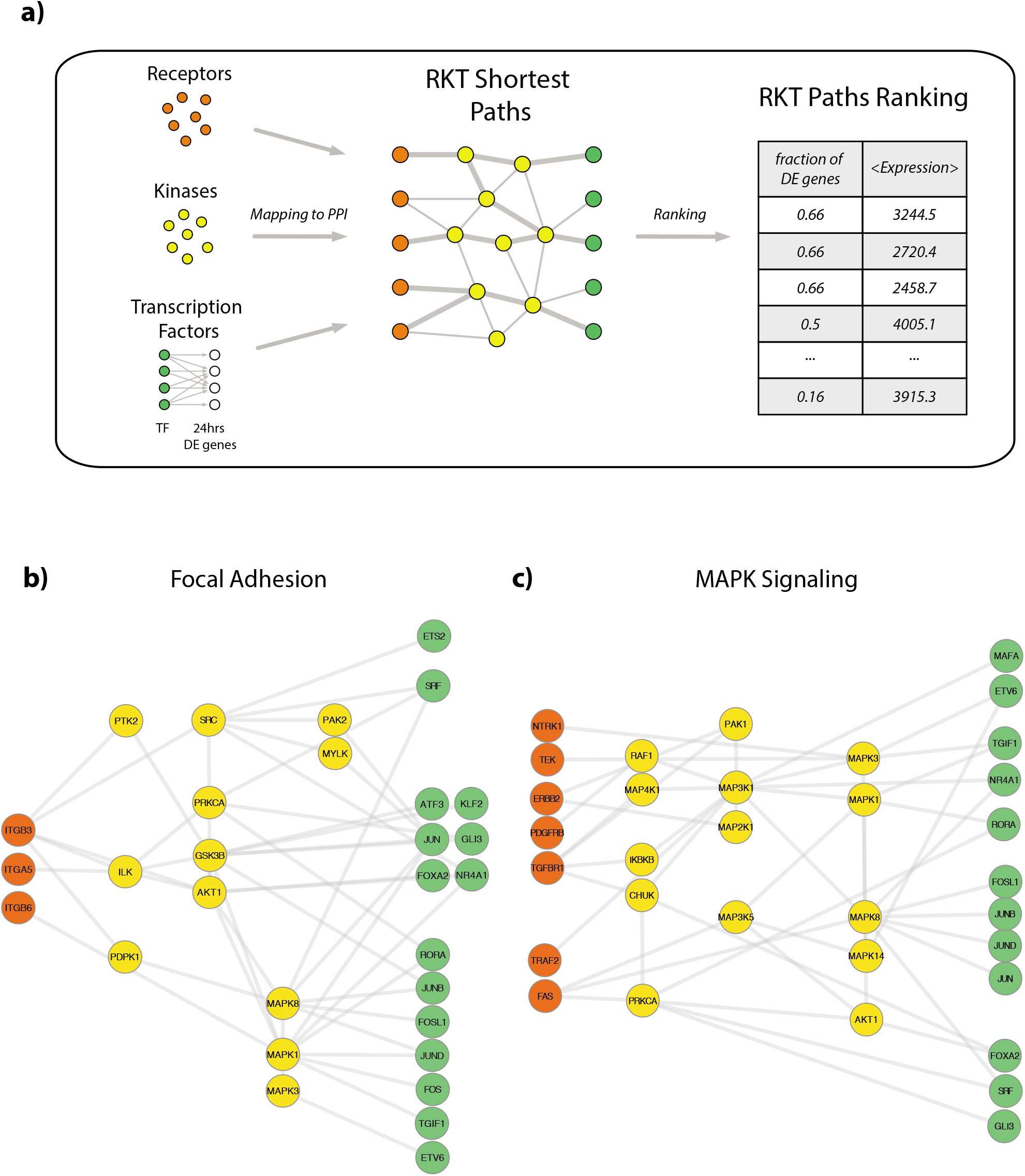
RKT Analysis. **a** Overview of the network analysis used to identify the RKT paths. First, the Protein-Protein Interaction (PPI) network is filtered for receptors, kinases, and transcription factors that impact DE genes at 24 hours. Within this subnetwork, we identified all the shortest paths (RKT paths) starting from a receptor, connecting up to 5 layers of kinases, and ending in a transcription factor. RKT paths were ranked based on the fraction of their nodes that are DE at 3 hours and on the average expression of their nodes at 3 hours. **b** Top-ranked RKT paths containing nodes annotated to the focal adhesion pathway. **c** Topranked RKT paths containing nodes annotated to the MAPK signaling pathway. It should be noted that PPI edges do not have directionality and, hence, the placement of the nodes in panels b and c has been chosen based on biological considerations associated with the RKT paths.

Starting from a recently proposed PPI^78^ (see Materials and Methods), we filtered the network to derive two subgraphs that respectively contained the receptors/kinases annotated to the focal adhesion and MAPK signaling pathway, as well as the set of transcription factors described above. We assumed a one-to-one correspondence between genes and their protein products in the entire analysis. For each signaling subgraph, we computed all the shortest paths connecting each receptor-TF pair (see Materials and Methods). We then ranked these RKT paths based on the fraction of DE genes in each path and used the average expression of the path’s nodes at 3 hours to break the ties (Fig. 4a). The top selected paths represent the most active downstream processes induced by compression (for a detailed list of all the network paths identified see Supplementary Tables 9a and 9b).

The top RKT paths within the focal adhesion subgraph were initiated by integrins ITGA and ITGB (Fig. 4b), showing a predominant role of these ECM receptors compared to receptor tyrosine kinases. The integrin-induced paths emphasized three main downstream signaling directions: 1) the engagement of *SRC* and *MYLK* kinases, both implicated in actin contractility^79,80^; 2) the activation of AP-1 transcription factors such as *JUN, JUNB*, and *ATF3* via the ILK-AKT1/GSK3B signaling path, known to regulate AP-1 activity through phosphorylation of *AKT1* and *GSK3B* by the kinase *ILK*^81^; 3) the activation of multiple *FOS* and *JUN* homodimers via *MAPK1* and *MAPK8*, highlighting the role of ERK2 and JNK signaling in the transmission of the mechanical stimulus.

A detailed view of the MAPK signaling RKT paths (Fig. 4c) revealed the engagement of various growth factor receptors, including *NTRK1, TGFB1R, TEK*, and *ERBB2*, as well as cytokine receptors (*FAS*). In agreement with the results in the focal adhesion subnetwork, the active RKT paths in the MAPK subgraph suggested the downstream regulation of the ERK1/2 pathway via the cascade *MAP3K1, MAPK1*, and *MAPK3* and of JNK pathway via *MAPK8*. Additionally, overrepresentation of RKT paths including *PRKCA, CHUK, IKBKB*, and the nuclear receptors *NR4A1* and *RORA*, highlighted the participation of the NFκB pathway in the signal transmission.

The focal adhesion and MAPK RKT paths pointed to the downstream engagement of 16 TFs. To determine the main processes regulated by these TFs, we performed pathway overrepresentation test of the 24 hours DE genes that are targeted by these 16 TFs (see Materials and Methods). Top GO Biological Processes (Supplementary Table 10 and Supplementary Fig. 2) included extracellular structure organization (p-adj.=1.3e-08), response to transforming growth factor beta (p-adj.=2.6e-08), and positive regulation of cell migration (2.6e-06). Notably, we found a large overrepresentation of metabolic processes involving biosynthesis of cholesterol and alcohol, supporting the activation of specific metabolic programs during UJT.

The RKT analysis predicted the initiation of signaling pathways that are consistent with the morphological changes observed during UJT. Reorganization of the actomyosin network and activation of AP-1 transcription factors are both essential mechanisms to regulate cellular shape and motility. Since these processes involve multiple components that are not only receptors, kinases, or transcription factors, we further investigated our network analysis considering the entire PPI.

### Flow centrality predicts key genes mediating the transcriptional response of HBECs to compression

We expanded our network analysis to determine, at a genome-wide level, which proteins had a major role in mediating the molecular processes triggered by compression. To achieve this goal, we considered a recently proposed ‘betweenness’ centrality measure called flow centrality^82^.

In network theory, the betweenness centrality of a node is defined as its frequency among all the shortest paths that traverse a network. Flow centrality extends this definition, encompassing only the shortest paths connecting any gene in a specific source subset to any gene in a specific target subset. The output of the procedure is a Flow Centrality Score (FCS), indicating the significance of the flow centrality of each node by comparing its value to a null distribution obtained by randomizing source and target gene sets (see Materials and Methods). Genes with high FCS, called “flow central genes”, are more likely to be involved in the interactions connecting source and target sets of interest, constituting a topological bottleneck in the communication between the two subsets.

In our study, we chose the DE genes at 3 hours and 24 hours respectively as source and target sets, and applied flow centrality to select the genes involved in the interactions between these transcriptional waves. A total of 556 flow central genes with FCS>2 was identified (see Material and Methods and Supplementary Table 11).

To determine potential functional similarities among these genes, we clustered them based on their shared source and target nodes and investigated the network properties of each of the resulting clusters. For each flow central cluster, we calculated a “connectivity score” 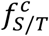. This quantity is defined as the average fraction of flow central genes connected to at least one source or target node (see Materials and Methods). High values of 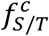 imply high connectivity of the flow central cluster with respect to the source/target set.

Our clustering analysis identified five main clusters (Fig. 5a and Supplementary Table 12). Interestingly, except for one cluster which had low connectivity to both source and target nodes, the identified four clusters exhibited strikingly different connectivity features with respect to the source and target genes. Cluster 1 and cluster 4 were highly connected to the source and target set, respectively (Fig. 5b). In contrast, cluster 2 and cluster 3 exhibited 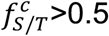 both for the source and target genes.

**Fig. 5.**
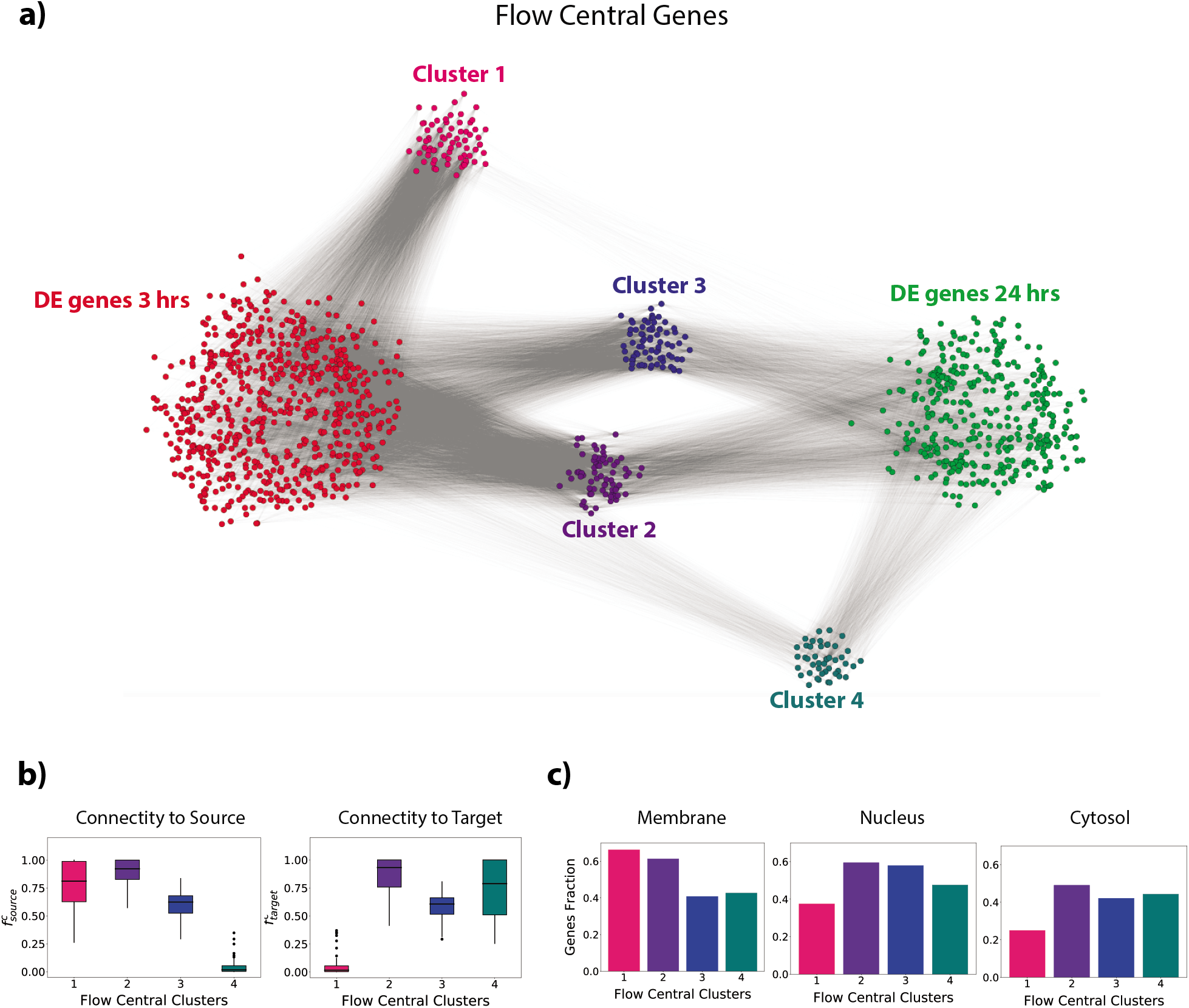
Flow Centrality Analysis. **a** Network visualization of the flow central clusters identified using flow centrality. **b** Connectivity score of each flow central cluster with respect to the source (left) and the target (right) gene sets. **c** Relative fraction of genes in each cluster that are annotated to the membrane, nucleus, and cytosol GO terms.

The connectivity score is proportional to the number of shortest paths connecting a cluster to the source/target and hence, indirectly provides information on the functional affinity of the clusters to the DE genes at the two time points. Since the DE genes at 3 hours were mainly associated with membrane proteins and DE genes at 24 hours were mainly associated with extracellular matrix and collagen proteins, the decreasing source connectivity and increasing target connectivity from cluster 1 to 4 suggested a gradual propagation of the signal from the membrane to the cellular nucleus, transmitted via the flow central clusters. To test this hypothesis, we looked at the relative abundance of each cluster in three main GO cellular components: the membrane (GO:0016020), the cytosol (GO:0005829), and the nucleus (GO:0005634). The fraction of genes localized in the membrane decreased from cluster 1 to 4 as source connectivity decreases, while proteins localized in the cytosol and nucleus correspondingly increased (Fig. 5c). This suggests that the flow central genes in cluster 1 represented the first mediators of the signal while the remaining clusters represented the gradual signal transduction to the proteins responsible for the late transcriptional response observed at 24 hours.

The list of flow central genes was consistent with the signaling pathways highlighted in the previous section. We compared the FCS distribution of the genes annotated to focal adhesion and MAPK signaling pathways to the FCS distribution of the remaining genes in the PPI. For both pathways, the FCS of annotated genes were statistically higher (Mann-Whitney U test p-val. = 1.3e-15 for focal adhesion and 6e-09 for MAPK signaling) than the FCS of other genes (Supplementary Fig. 3). It follows that flow centrality supported a leading role of focal adhesion molecules and MAP kinases in the communication between the two transcriptional responses at 3 hours and 24 hours.

Flow central nodes included multiple genes identified in the RKT analysis. Flow central genes annotated to either the focal adhesion or MAPK signaling pathway overlapped significantly (hypergeometric p-val.<1e-03) with the genes of the RKT paths, such as *PTK2, AKT1, MAPK1*, and *MAPK8*, and included additional components, thus confirming and extending by an independent methodology the network predictions obtained through the RKT path analysis.

Several integrins showed high FCS, including *ITGAV, ITGB3, ITGB4, ITGB5*, and *ITGB6* (Fig. 6a). These molecules bind to ECM proteins with their extracellular head region and to the actin cytoskeleton via intracellular proteins that attach to their cytoplasmic domain^83^. In support of this mechanism, both matrix proteins *SPP1* and *FN1*, and adapter proteins *FLNB* and *FLNA*, displayed high FCS (Supplementary Table 11). Additional flow central genes included *PTK2, PAK4*, and *CAPN2*, central bottlenecks for the regulation of actin polymerization and for propagation of downstream signaling pathways within the cell^84,85^.

**Fig. 6.**
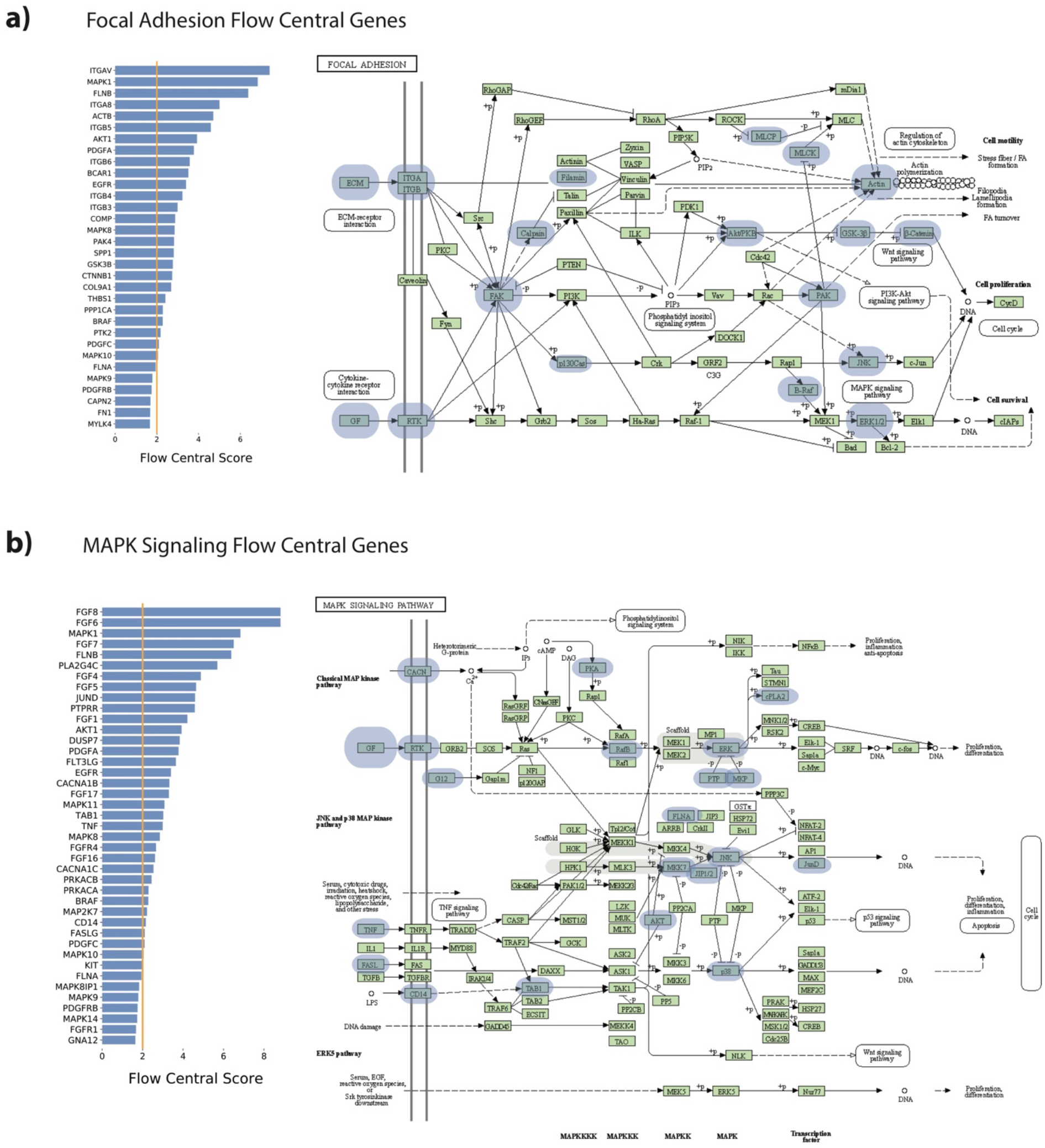
Flow central genes associated with the a) focal adhesion and b) MAPK signaling pathways. For each pathway, the flow central score (FCS) of the top flow central genes is shown in the left panel. In the right panel, the location of each of the flow central genes within the KEGG pathway representation is highlighted.

Among these pathways, we found high FCS of the JNK-associated components *MAP2K7, MAPK8/9/10, MAPK8IP1*, and *JUND*, as well as to the ERK2-associated proteins *BRAF* and *MAPK1*, which showed a high FCS of 6.83 (Fig. 6b). The overrepresentation of genes involved in the ERK and JNK signaling pathways corroborated the crucial contribution of these processes to mechanical stimulus transduction in HBE cells. Additionally, the high FCS of *AKT1, GSK3B*, and *CTNNB1*, known to modulate cellular apoptosis via regulation of β-catenin activity^86^ (Fig. 6a), provided a possible explanation for the inhibition of cell proliferation previously observed in our system^11,15^.

We also observed high FCS of multiple fibroblast growth factors and their corresponding receptors. This result may imply a parallel activation of ERK pathway via signal transmission to adaptor proteins like *GRB2, SOS*, and *Ras*^87^, as well as hint at a potential cross-talk between these receptors and integrins, as already reported in ^88^.

Overall, our analysis showed that the morphological changes of UJT are characterized at the molecular level by actin repolymerization, JNK- and ERK-mediated regulation of cell motility, and β-catenin reorganization. This suggests that cells undergoing UJT trigger a specific transcriptional program that allow remodeling of the cytoskeleton network and of the adhesive interactions with the ECM.

### Flow centrality identifies ECM remodeling proteins as key regulators of the structural changes induced by compression on asthmatic HBE cells

Park et al.^11^ showed that unjamming is not only relevant in the context of mechanotransduction but that it also has implications for aberrant airway remodeling that is a cardinal feature of asthma. Compared with HBECs derived from non-asthmatic donors, HBECs derived from asthmatic donors exhibit phenotypic characteristics of unjammed tissues, with more elongated shapes and prominent collective migratory behavior^11^. At the transcriptional level, a recent study has shown that asthmatic HBECs and nonasthmatic HBECs at 24 hours post-compression exhibit similar transcriptional patterns as genes associated with airway remodeling and ECM reorganization^30^.

Similarities between asthmatic HBECs at baseline and non-asthmatic HBECs post-compression can be traced back to the effect of asthmatic bronchoconstriction on the airway. During bronchoconstriction, the airway becomes extensively narrowed and contracted, and airway caliber is reduced^89^, resulting in excessive mechanical compression of the epithelial layer^90^ (similar to ~30 cm H2O compression). A similar compression applied in our experimental system recapitulates key anatomic and pathophysiologic features as shown during asthmatic bronchoconstriction^23,24,26,29,37,91^, so we investigated the potential existence of common molecular components involved in the unjamming of both asthmatic and compressed normal epithelial cells. To achieve this task, we assessed the transcriptional profile of ALI-cultured HBECs from four asthmatic donors through bulk RNA-Seq. RNA-Seq data were extracted at baseline consistent with the experimental setup of compressed normal HBECs (see Materials and Methods). We performed differential expression analysis on the asthmatic HBECs versus the control, unperturbed HBECs at baseline using the R package DESeq2. By choosing a threshold of FC>1.5 and FDR-adjusted p-value < 0.05, we identified 253 DE genes (see Materials and Methods and Supplementary Table 13).

We compared DE genes in non-asthmatic HBECs at 24 hours post-compression to DE genes in asthmatic HBECs at baseline (no pressure) using the same FC threshold (FC>1.5). The two sets overlapped significantly (hypergeometric p-val.=4.6e-12) and the sign of the FC of the DE genes at 24 hours post-compression and in asthmatic HBECs were positively correlated (correlation coefficient *ρ* = 0.4, p-val=1.8e-09) (see Materials and Methods and Supplementary Fig. 4). Next, we applied flow centrality. We selected the DE genes at 3 hours post-compression in normal HBECs as the source gene set, and we considered two different sets of genes as targets: (1) the DE genes at 24 hours post-compression in normal HBECs and the (2) DE genes in asthmatic cells without any perturbation. In order to remove potential similarities due to the same source genes, the FCS were computed by randomizing only the target module (see Materials and Methods). The two resulting sets of FCS, displayed in Fig. 7, represented the specificity of each PPI node in connecting the source to the target modules as compared to a random set of target genes.

**Fig. 7.**
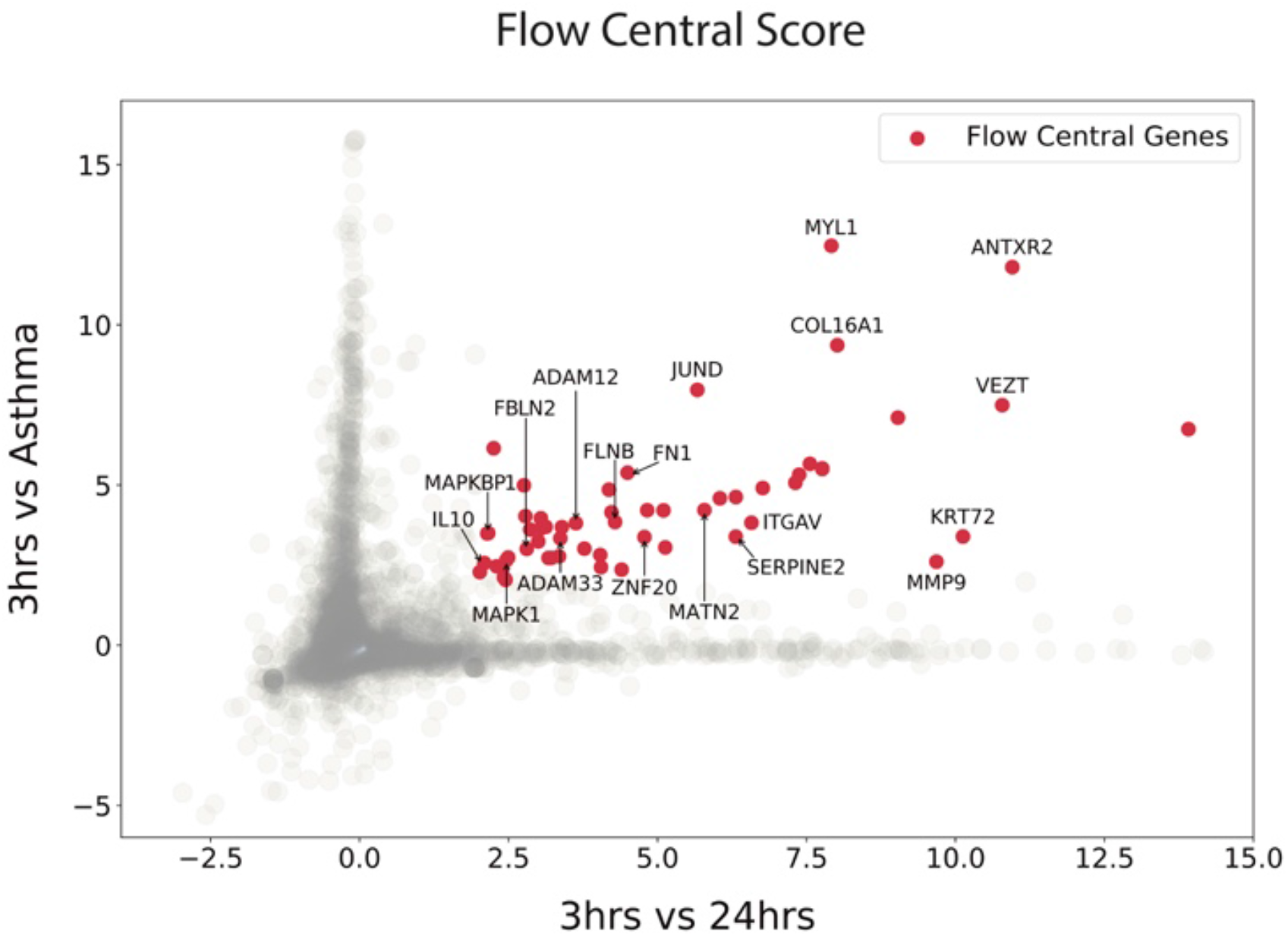
Flow centrality in asthma. The FCS of each PPI node calculated by applying flow centrality to genes DE at 3 hours versus genes DE at 24 hours (x-axis), as well as genes DE at 3 hours versus genes DE in asthma (y-axis). Common flow central genes (FCS>2 in both analyses) are highlighted in red.

While multiple genes showed different values of FCS between the two datasets, a statistically significant subset of 57 flow central genes (Supplementary Table 14) appeared to mediate both the transcriptional response of non-asthmatic HBECs at 24 hours post-compression and asthmatic HBECs at baseline. Many of these common flow central genes are implicated in the regulation of ECM, cell-matrix interactions, and cell-cell adhesive junctions. High values of the FCS were assigned to both ECM proteins such as *COL16A1, FN1, MATN2*, and *FBLN2* and to focal adhesion proteins such as *ITGAV* and *FLNB*. Common flow central genes also included *MAPKBP1, MAPK1*, and *JUND*, highlighting the role of JNK and ERK signaling cascades in both phenomena. In both asthmatic and compressed normal HBECs, flow centrality identified molecules like *VEZT*, a component of the cadherin-catenin complex critical to the formation of adherens junctions^92^, and the metalloproteinases *ADAM12* and *MMP9*, proteins known to be involved in the degradation of the ECM^93,94^, as key mediators of the signal transmission. Not only are these genes involved in the structural reorganization of the intercellular and ECM-interactions^95,96^, they have also been shown to be associated with the airway remodeling processes observed in asthmatic airways as a consequence of bronchoconstriction^27^.

Our analysis linked, at the genomic level, the response of non-asthmatic HBEC to pressure both to the UJT and the tissue remodeling in asthmatic airways. We identified common genomic patterns that induce ECM remodeling in both asthmatic and non-asthmatic unjammed HBECs, thus suggesting at the molecular level a connection between airway remodeling, bronchoconstriction and UJT as described in previous studies^21,26,27,29,36,37,91,97^

## Discussion

Our analysis shows that normal HBECs respond to compression in a multiphasic manner. Immediately after pressure release (3-hour time point), HBECs transcriptionally reprogram cytoskeletal genes regulating the structure of the actomyosin network; at 21 hours after pressure release (24-hour time point), HBECs activate a delayed transcriptional response involving the ECM and its interaction with cellular basal membrane. By using a network-based approach, we show that these genomic patterns are the result of multiple, coordinated signaling pathways that induce cell shape changes via actin repolymerization processes and induce increased cell motility via ERK- and JNK-mediated activation of AP-1 transcription factors.

The RNA-Seq data and network analysis highlighted the central role of integrins. Integrins are heterodimeric proteins consisting of an alpha and beta chain that can be combined to give rise to 24 unique integrin molecules^83^. By binding to cytoskeletal and matrix proteins, these transmembrane receptors serve both as signaling centers and traction points during cell migration^98^. Force transmission mechanisms mediated by integrins have been extensively described over the past decade^14,99,100^. Nevertheless, limited knowledge has been achieved on the systematic relationship between these receptors and cellular motions. Our study revealed that integrins were differentially expressed throughout the entire process of unjamming (up to 24 hours after initial pressure exposure) and that they were involved in multiple signaling pathways. This result characterized integrins as more than simple mechanoreceptors and hinted at the potential activation of a positive feedback loop where integrins steadily recruit various molecular components to assist and stabilize cellular unjamming.

In agreement with this scenario, our analysis showed that these receptors both induce actin polymerization processes via *SRC* and focal adhesion kinase *PTK2*, and initiate additional downstream signals to activate the transcription factors *JUN* and *FOS* via ERK1/2 and JNK pathways. Even though these pathways are known to be triggered in response to mechanical stress^23,101^, the picture that emerged from our study indicated that ERK1/2 and JNK are active regulators of cell motility at several hours since the initial application of pressure. Notably, both these signaling pathways have been previously associated with the collective migration of cells in epithelial sheets^102-108^. Propagation of ERK1/2 waves have been observed to be correlated with collective cell movements in a wound healing assay of MDCK cells^102,103,107^. The JNK pathway has been also connected to the regulation of collective migration of several cell types in wound healing and developmental processes^104-106,108,109^.

Our analysis takes advantage of transcriptomics and molecular interaction data to describe the UJT at the genome-wide level, yet it suffers from some limitations. First, the integration of additional types of -omics data and a larger sample size would allow us to further investigate undetected molecular processes such as epigenetic and post-translational modifications. Larger number of subjects would be crucial especially for the comparison with asthmatic cells, given the heterogeneity of this disease.

Moreover, the RNA isolation process prevents the epithelial tissue from being sequenced and analyzed through living imaging at the same time and in the same sample. We have had to rely on separate experiments under similar conditions to make the inferences described herein.

Mechano-transduction processes do not necessarily require differential expression of their involved genes and can be activated as early as seconds after an external stimulus^110-113^. It follows that the signaling pathways identified by our study likely represent the molecular processes sustaining the UJT, leaving room for speculation on what is the initial trigger that induces this phenomenon. Previous studies performed by our, and other, groups have shown that mechanical compression on HBECs stimulates downstream ERK signaling by activating *EGFR* and its ligands only 20 minutes after pressure application, suggesting the possible pre-existence of latent forms of these growth factors in the epithelial tissue^21^. Additionally, our RNA-Seq data are compatible with the possible activation of cell receptors via conformational changes or physical aggregation of these molecules, irrespective of their specific ligand. For example, compression and stretch can directly induce structural changes in integrins^114-116^, growth factor receptors^117-120^, intracellular adaptor kinases^121-125^ as well as ion channels^110,126^. These alterations have been reported together with rapid activation of downstream intracellular pathways, including PI3K, ERK, and JNK signaling^110,114-126^. Given the pronounced transcriptional response we observed at the 3-hour time point, it is conceivable that similar events occur *in vivo* in compressed airway epithelial cells during asthmatic bronchospasm.

These findings, taken together, show that the UJT program is not the result of any single signaling pathway but rather is a highly complex, dynamic, and non-linear process that involves a coordinated interplay of downstream pathways including development, fate selection, energy metabolism, cytoskeletal reorganization, and extracellular matrix remodeling.

## Author contributions

M.D.M., A.K. and E.M. analyzed data. M.D.M., A.K., K.G. and S.T.W. interpreted data. M.D.M. and A.K. wrote the manuscript. J.M., M.M., R.C., J.J.F., J.A.P. and S.T.W. designed the experiment and generated the experimental data. All the authors contributed to the writing of the paper, provided critical feedback and helped shape the research and the analysis of the problem.

## Acknowledgements

This work was funded by the National Cancer Institute (NCI grant number U01CA202123), the National Heart Lung and Blood Institute (NHLBI grant numbers NHLBI 1R01HL148152, UH3 OD023268, P01HL120839, P01HL132825, T32 HL007118, and K25HL133599), the Parker B. Francis Foundation, the Lemann Foundation, the German National Science Foundation (Deutsche Forschungsgemeinschaft), and the American Heart Association (13SDG14320004). This work was conducted with support from Harvard Catalyst | The Harvard Clinical and Translational Science Center (National Center for Advancing Translational Sciences, National Institutes of Health Award UL 1TR002541) and financial contributions from Harvard University and its affiliated academic healthcare centers. The content is solely the responsibility of the authors and does not necessarily represent the official views of Harvard Catalyst, Harvard University and its affiliated academic healthcare centers, or the National Institutes of Health.

## Materials and Methods

### Primary human bronchial epithelial cells and mechanical compression

Primary Human Bronchial Epithelial Cells were derived from 4 donors with no pre-existing chronic lung disease and 4 asthmatic donors following the same protocol described in ^11^. HBECs were grown for 5-6 days in order to reach confluency and subsequently cultured in air-liquid interface (ALI) conditions^11^. For each donor, two independent replicates were introduced into the polarization process.

On ALI day 14, well-differentiated non-asthmatic HBECs were exposed to 30 cm H_2_O apical-to-basal mechanical compression for 3 hours as described in ^11^ and after this time pressure was released. Compressed non-asthmatic HBECs were harvested both immediately after pressure release, referred in the text as the 3-hour time point, as well as after a further incubation period of 21 hours, referred in the text as the 24-hour time point. Control and asthmatic non-compressed HBECs were also harvested at the same reference time points. Harvested cells were used for isolation of RNA.

### RNA-isolation, library preparation and RNA sequencing

Total RNA was isolated organically using QIAzol lysis reagent and the Qiagen miRNeasy Kit (Qiagen). Quality was assessed using the Nanodrop 8000 spectrophotometer. Sequencing libraries were constructed with the TruSeq^®^ Stranded Total RNA Library Prep Globin Kit (Illumina). Sequencing was performed using a HiSeq 2500 instrument (Illumina). Raw reads were trimmed with Skewer^127^ and then mapped to the GRCh38 reference genome using STAR^128^. Read counts were computed with HTSeq^129^. Data were normalized by using the DESeq2^130^.

### Differential expression analysis

Differential expression analysis was performed using the R package DESeq2^130^ (v.1.22.2). Three separate DE analyses were performed: non-asthmatic compressed HBECs at 3 hours versus non-asthmatic uncompressed HBECs at 3 hours, non-asthmatic compressed HBECs at 24 hours versus non-asthmatic uncompressed HBECs at 24 hours, and asthmatic versus non-asthmatic uncompressed HBECs. Genes with zero counts across all the different conditions were removed. For the first two DE analysis, the design matrix was built to take into account batch effects in cells derived from the same donor. In the third DE analysis between uncompressed asthmatic versus non-asthmatic HBECs, both the 3-hour and 24-hour time points were included. The design matrix was implemented to remove temporal batch effects. In all the DE analyses, we applied DESeq2 independent filtering and p-values were corrected for multiple hypothesis testing by using the Benjamini-Hochberg (BH) false discovery rate (FDR) adjustment^131^.

### Hierarchical clustering to identify the three transcriptional regimes

The three transcriptional regimes (transient, steadily-increasing, and long-term) described in the main text were identified by performing hierarchical clustering on the z-scores of the normalized expression counts. Specifically, raw counts were normalized and transformed for variance stabilization using the DESeq2 function VST. We then performed a hierarchical clustering on the counts’ z-scores, using the Euclidean distance and the complete-linkage metrics. We cut the cluster dendrogram so that we obtained 2 clusters for genes DE only at 3 hours, 6 clusters for genes DE only at 24 hours, and 4 clusters for genes DE at both 3 hours and 24 hours. Based on the clustering results, we selected three specific clusters corresponding to three characteristic transcriptional responses: transient, long-term and steadily increasing.

### Enrichment analysis: Gene Ontology, KEGG, and Reactome enrichment analysis

For each transcriptional regime, we performed Gene Ontology overrepresentation tests using the “enrichGO” function in the R package “clusterProfiler”^43^ (v.3.10.1) with default settings. In both GO Cellular Components and Biological Processes terms, we used the R annotation package “org.Hs.eg.db” (v.3.7.0) and the BH-FDR method for p-value adjustment.

In order to identify the most active pathways mediating the different transcriptional regimes, we used the clusterProfiler function “enrichKEGG” to perform KEGG pathway enrichment on the merged set of DE genes at 3 hours and 24 hours. P-values were adjusted using the BH-FDR correction.

We confirmed the results of the KEGG enrichment using also the Reactome database^67^. We performed enrichment analysis on the merged set of DE genes at 3 hours and 24 hours using the clusterProfiler function “enrichPathway”. P-values were adjusted using the BH-FDR correction and we selected pathways enriched with p-adj.<0.05.

### Protein-protein interaction network

The protein-protein interaction (PPI) network used in our network analysis was compiled by Cheng et al.^78^. As described in the original paper, the PPI network integrates 15 different databases and include 1) binary PPIs identified via high-throughput yeast-two-hybrid (Y2H) experiments^32,132^, 2) kinase-substrate interactions from literature-derived low-throughput and high-throughput experiments^133-139^, 3) literature-curated PPIs identified through mass spectrometry (AP-MS), Y2H, literature-derived low-throughput experiments, and protein three-dimensional structures^140-145^, and 4) signaling network interactions from literature-derived low-throughput experiments, as annotated in SignaLink2.0^146^. Only the largest connected component of the network was considered, resulting in an interactome of 16,656 proteins and 243,592 interactions.

### RKT analysis

As an initial input for the implementation of the Receptor-Kinase-Transcription Factor (RKT) analysis, we filtered the PPI network based on a list of receptors (R), kinases (K), and transcription factors (TFs) which were identified as follows. Receptors and kinases were assembled from the curated databases developed in Ref. ^76^ and Ref. ^77^ respectively. For the transcription factors, we first generated a list of motif mappings. To do this, we used FIMO^147^ to scan the human genome (hg38) for a comprehensive set of CIS-BP Single Species DNA-motifs curated in the MEME suite^148^. Hits that met a significance more than 1e-4 and fell within [-750, +250] base pairs of a gene’s transcriptional start site (based on gene annotations downloaded from UCSC^149^) were used to construct a set of TF-gene interactions. We used this regulatory network to select transcription factors targeting genes differentially expressed at 24 hours post-compression. In particular, for each TF, we used the hypergeometric test to compute the p-value of the overlap between the TF’s target genes and DE genes at 24 hours. TFs with a p-value lower than 0.05 were included in the network.

After filtering the PPI network based on this list of receptors, kinases, and transcription factors, we removed all the R-R, R-TF, K-R, T-R, T-K, T-T edges in order to restrict our analysis to the shortest paths following the RKT directionality. We further filtered the network to include R and K annotated specifically to the KEGG Focal Adhesion (hsa04510) and MAPK (hsa04010) pathways, obtaining two separate subgraphs for each pathway. For each subgraph, we computed all the shortest paths containing one layer of receptor, up to five layers of kinases and one layer of transcription factors. Paths that included interactions between different receptor classes were removed, as these interactions only occur indirectly via crosstalk of downstream signaling pathways and do not represent direct mechanistic interactions of the receptor chains.

To determine the main processes regulated by the most active RKT paths, we selected the genes that were targeted by TFs in each RKT path based on the regulatory network described above and that were DE at the 24-hour time point. We performed pathway overrepresentation test on the GO Biological Processes enriched in this gene set using clusterProfiler (see above) and we focused on the top 30 enriched pathways (BH-FDR p-adj.<0.05). We used all of the target genes (DE and not-DE) of TFS in the RKT paths as background for the overrepresentation test.

### Flow Centrality and Flow Connectivity Score

Flow centrality is a topological measure proposed by Maiorino et al.^82^. The flow centrality *FC^S,T^*(*i*) of a node *i* with respect to a given source *S* and target *T* set of nodes is given by:

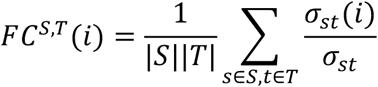

where *σ_st_*(*i*) is the number of shortest paths connecting a source node *s* to a target node *t* passing through node *i, σ_st_* is the total number of shortest paths connecting *s* and *t*, and |*S*| and |*T*| denote the size of the source and target sets, respectively. Given this definition, the flow central score (FCS) of a node represents the statistical significance of a node’s flow centrality value with respect to a null distribution of random pairs of source and target gene sets. Specifically, the FCS of a node *i* is equal to:

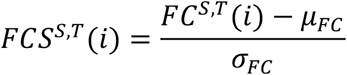

where *μ_FC_* and *σ_FC_* are the average and standard deviation across the distribution of the node’s *FC^S,T^*(*i*) values obtained by computing its flow centrality with respect to 1000 random source and target gene sets (see ^82^ for details). High values of FCS indicate that the node occurs more frequently in connecting the source and target modules. Genes with FCS higher than 2 in the PPI are defined as “flow central genes”. In contrast to the raw values of flow centrality, the flow central score is not biased toward high degree nodes.

In order to determine potential similarities among the 556 flow central genes identified in our analysis, we performed a clustering analysis. For each flow central gene, we created a binary vector representing all source and target nodes; we assigned a value of 1 to an element of this vector if the corresponding source/target node occurred in one of the shortest paths mediated by the flow central gene. We performed an unweighted average linkage clustering on these 556 binary vectors using the Hamming distance metric and identified 5 main clusters. We then investigated the connectivity properties of each cluster by defining a connectivity score, 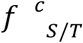, as:

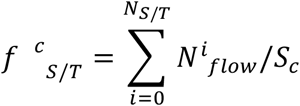

where 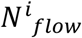 is the number of flow central genes in the cluster *c* that connect the source/target node *i, S_c_* is the size of the flow central cluster *c*, and *N_S/T_* is the size of the source and target modules respectively. The connectivity score measures the relative amount a cluster mediates the communication with the source/target sets.

### Comparison between flow central genes in asthmatic HBECs at baseline and in non-asthmatic HBECs at 24 hours post-compression

We first compared the transcriptional profiles of 24 hours post-compression non-asthmatic HBECs and non-compressed asthmatic HBECs by computing the overlap DE genes in each condition. We found 14 significantly overlapping genes between DE genes at 24 hours post-compression and DE genes in asthmatic cells (hypergeometric p-val.= 4.64e-12). To further identify global patterns of similarities, we selected the DE genes at 24 hours post-compression (FC>1.5 and BH-FDR p-adj.<0.05) and we compared the sign of their FCs with respect to asthmatic cells. The two sets of FCs resulted positively correlated with Pearson correlation coefficient ρ=0.4 and two-tailed p-value=1.8e-09, showing similar perturbations in the genomic patterns of non-asthmatic post-compression cells and asthmatic cells.

For the comparison between the flow central genes connecting non-asthmatic post-compression HBECs at 3 hours versus 24 hours and non-asthmatic post-compression HBECs at 3 hours versus asthmatic HBECs, we modified the original definition of FCS to remove non-informative similarities in the calculation of the flow centrality values due to the same source set. To account for these biases, we randomized only the target module in the computation of the FCS, following the same randomization protocol described in ^82^. It follows that the resulting FCS expresses the statistical significance of the flow central values by comparing them with a null distribution where only the target set is variable. This approach allowed us to estimate similarities between flow central genes that are related only to the specific target sets chosen.

