## Supplementary Figures for "Genomic signatures of the unjamming transition in compressed human bronchial epithelial cells"

### Supplementary Figure 1

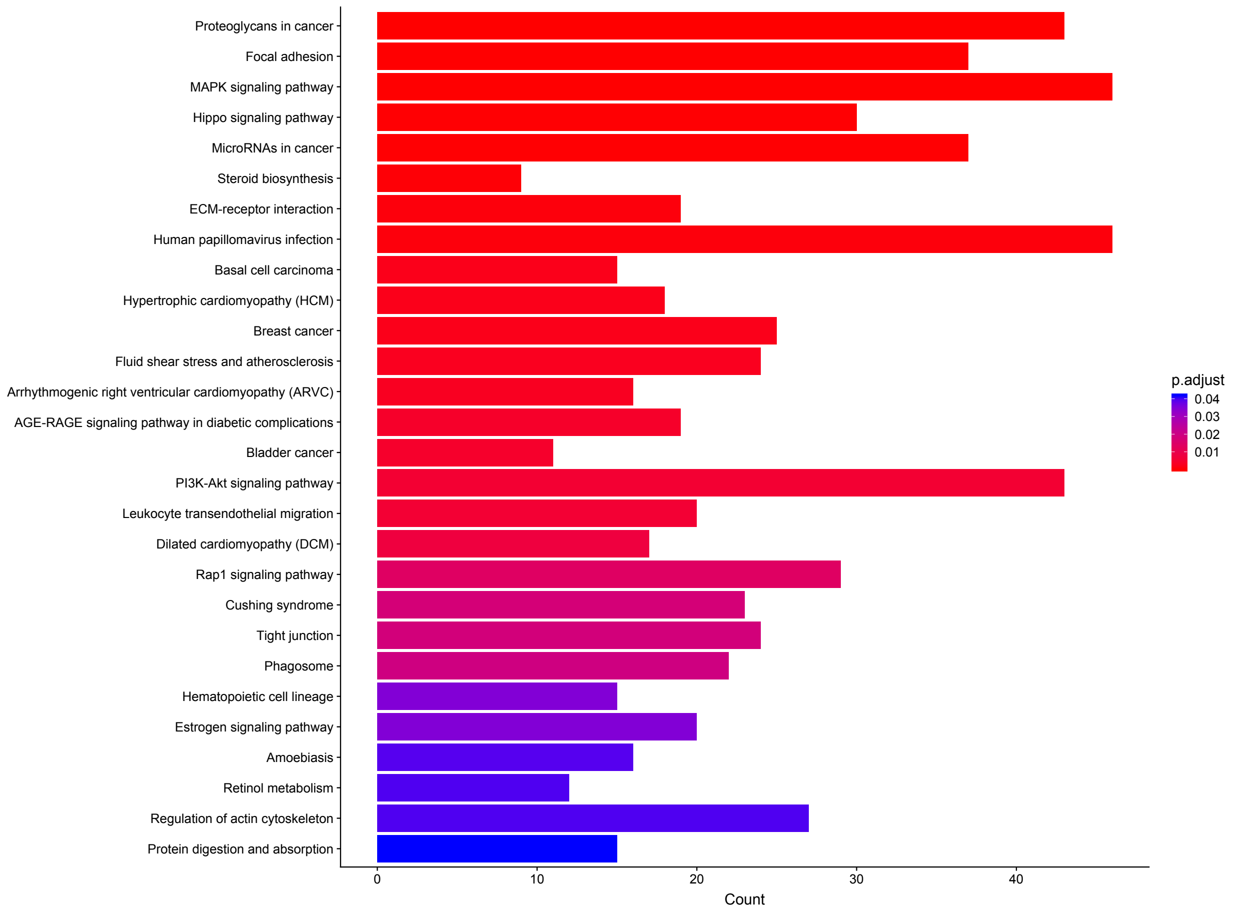


**Supplementary Figure 1.** List of KEGG pathways enriched (p-adj. < 0.05) in the merged set of DE genes at 3 hours and 24 hours post-pressure. The x-axis of the bar plot represents the number of genes annotated to the pathway and the color represents the value of the FDR-adjust p-value based on the colormap on the right side of the figure.

### Supplementary Figure 2

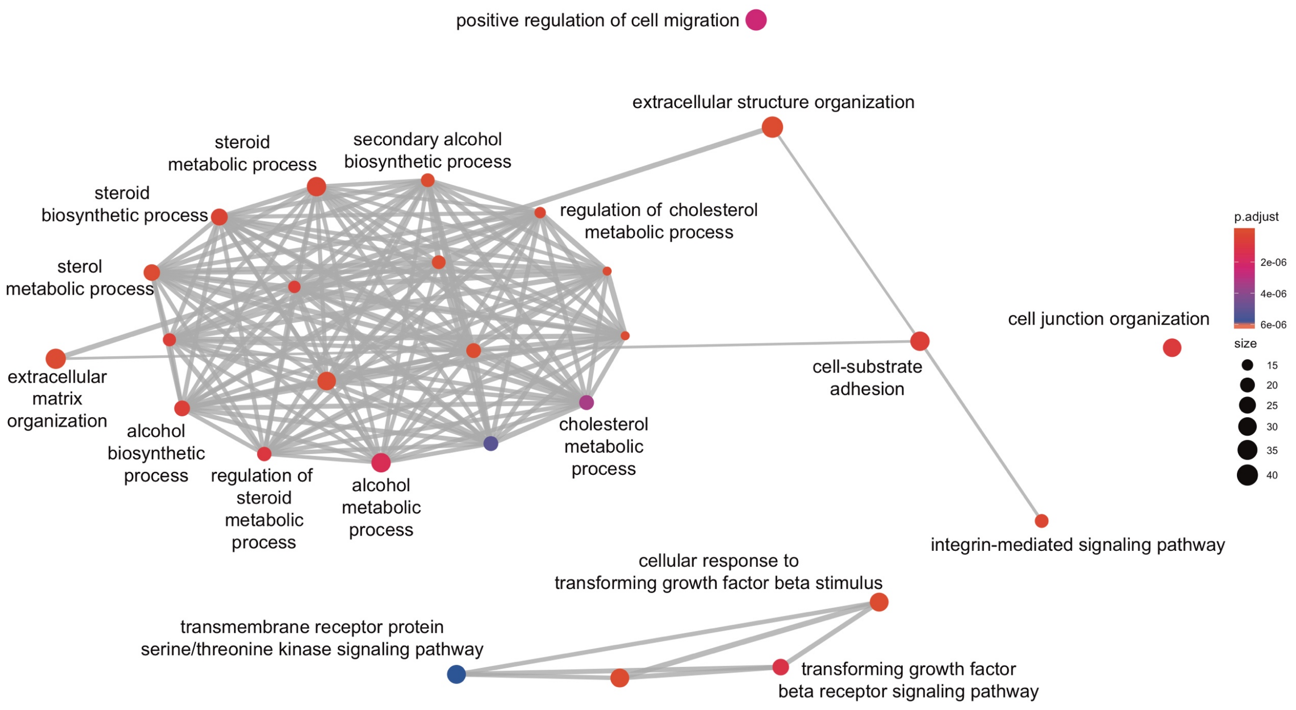


**Supplementary Figure 2.** Top 30 GO Biological Processes (BP) enriched in the subset of genes that are DE at 24 hours post-pressure and are targeted by the transcription factors identified in the RKT analysis. We visualized the BP enriched using a network representation where each node represents a pathway and edges between pairs of nodes represent common DE genes annotated to both pathways. For visualization purposes we showed only edges with more than 3 common DE genes. Node size and color are based on the number of DE genes annotated to the pathway and its adjusted p-value, respectively.

### Supplementary Figure 3

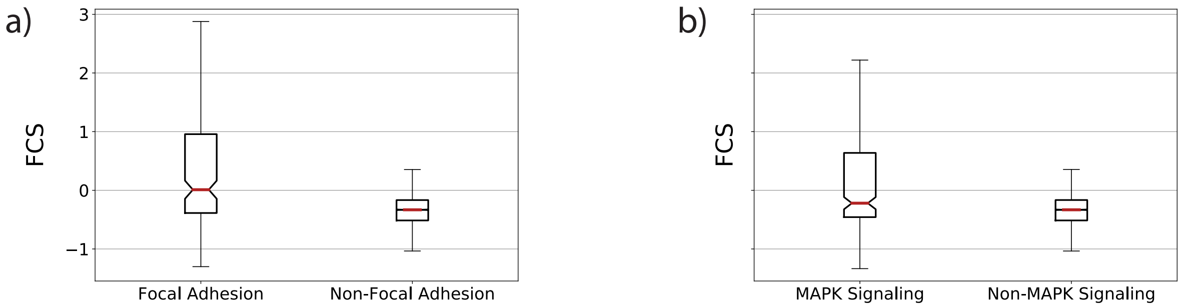


**Supplementary Figure 3.** Notched box plot of the FCS of **a** genes annotated to KEGG focal adhesion pathway compared to the remaining genes on the PPI, **b** genes annotated to KEGG MAPK signaling pathway compared to the remaining genes on the PPI.

Supplementary Figure 4

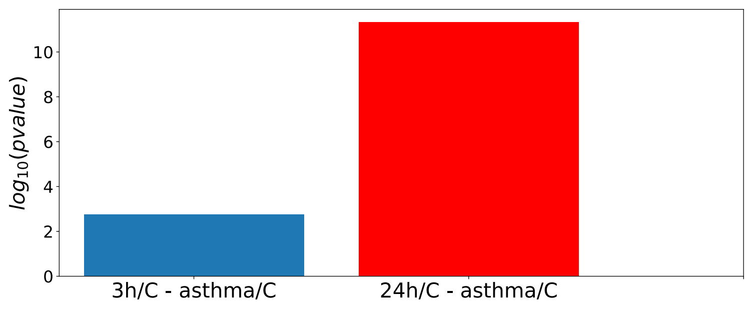


**Supplementary Figure 4.** Hypergeometric p-value of the overlap between genes DE at 3 hours (blue) and 24 hours (red) post-pressure versus genes DE in asthmatic HBECs at baseline.
