## Supplementary Tables Legends for "Genomic signatures of the unjamming transition in compressed human bronchial epithelial cells"

**Supplementary Table 1.** List of DE genes at the 3 hour time point obtained by performing differential expression analysis of compressed HBECs after 3 hours post-pressure with respect to uncompressed HBECs at the same time point. For each gene, we included its Ensembl ID, log-fold-change, FDR-adjusted p-value, and gene symbol.

**Supplementary Table 2.** List of DE genes at the 24 hour time point obtained by performing differential expression analysis of compressed HBECs after 24 hours post-pressure with respect to uncompressed HBECs at the same time point. For each gene, we included its Ensembl ID, log-fold-change, FDR-adjusted p-value, and gene symbol.

**Supplementary Table 3. a** List of DE genes that show the *transient upregulated* transcriptional response described in the main text. For each gene, we included its Ensembl ID, gene symbol, average normalized expression counts at baseline condition (average over 3 hours and 24 hours), at 3 hours post-pressure and at 24 hours post-pressure. **b** List of DE genes that show the *transient downregulated* transcriptional response described in the main text. For each gene, we included its Ensembl ID, gene symbol, average normalized expression counts at baseline condition (average over 3 hours and 24 hours), at 3 hours post-pressure and at 24 hours post-pressure.

**Supplementary Table 4.** List of DE genes that show the *long-term* transcriptional response described in the main text. For each gene, we included its Ensembl ID, gene symbol, average normalized expression counts at baseline condition (average over 3 hours and 24 hours), at 3 hours post-pressure and at 24 hours post-pressure.

**Supplementary Table 5.** List of DE genes that show the *steadily-increasing* transcriptional response described in the main text. For each gene, we included its Ensembl ID, gene symbol, average normalized expression counts at baseline condition (average over 3 hours and 24 hours), at 3 hours post-pressure and at 24 hours post-pressure.

**Supplementary Table 6.** List of GO Biological Processes enriched in: **a** the transient downregulated regime, **b** the transient upregulated regime, **c** the long-term regime, and **d** the steadily-increasing regime. For each pathway, we included its description, GO id, FDR-adjusted p-value and the set of genes annotated to the pathway.

**Supplementary Table 7.** Subset of GO Biological Processes that are enriched in the DE genes at 3 hours and are related to metabolic processes. For each pathway, we included its description, GO id, FDR-adjusted p-value and the set of genes annotated to the pathway.

**Supplementary Table 8.** Top 20 Reactome Pathways enriched in the merged set of DE genes at the 3 hour and 24 hour time point. For each pathway, we included its description, Reactome id, FDR-adjusted p-value and the set of genes annotated to the pathway.

**Supplementary Table 9.** Complete list of Receptors-Kinases-Transcription Factors (RKT) paths identified in **a** the focal adhesion subnetwork and **b** the MAPK signaling subnetwork.

**Supplementary Table 10.** Top 30 GO Biological Processes enriched in the subset of genes that are DE at 24 hours post-pressure and are targeted by the Transcription Factors identified in our RKT analysis. For each pathway, we included its description, GO id, FDR-adjusted p-value and the set of genes annotated to the pathway.

**Supplementary Table 11.** List of Flow Central genes mediating the signal between DE genes at 3 hours and DE genes at 24 hours post-pressure. For each gene, we included its gene symbol, flow central score (FCS), if it is included in the source gene set or if it is included in the target gene set.

**Supplementary Table 12.** List of genes belonging to each of the 5 flow central clusters identified. The identification number of each flow central cluster corresponds to the one used in the main text.

**Supplementary Table 13.** List of DE genes in asthmatic HBECs obtained by performing differential expression analysis of non-asthmatic HBECs at baseline versus asthmatic HBECs at baseline. For each gene, we included its Ensembl ID, log-fold-change, FDR-adjusted p-value, and gene symbol.

**Supplementary Table 14.** List of Flow Central genes that show FCS>2 choosing DE genes at 3 hours post-pressure as source set and either DE genes at 24 hours post-pressure (second column) or DE genes in asthmatic HBECs at baseline (third column) as target sets.
